# BrainLine: An Open Pipeline for Connectivity Analysis of Heterogeneous Whole-Brain Fluorescence Volumes

**DOI:** 10.1101/2023.02.28.530429

**Authors:** Thomas L. Athey, Matthew A. Wright, Marija Pavlovic, Vikram Chandrashekhar, Karl Deisseroth, Michael I. Miller, Joshua T. Vogelstein

## Abstract

Whole-brain fluorescence images require several stages of computational processing to fully reveal the neuron morphology and connectivity information they contain. However, these computational tools are rarely part of an integrated pipeline. Here we present BrainLine, an open-source pipeline that interfaces with existing software to provide registration, axon segmentation, soma detection, visualization and analysis of results. By implementing a feedback based training paradigm with BrainLine, we were able to use a single learning algorithm to accurately process a diverse set of whole-brain images generated by light-sheet microscopy. BrainLine is available as part of our Python package brainlit: http://brainlit.neurodata.io/.

## Main

Whole-brain image volumes at the micron scale are helping scientists characterize neuron-level morphology and connectivity, and discover new neuronal subtypes. These volumes require intense computational processing to uncover the rich neuronal information they contain. Currently, however, image acquisition is outstripping the availability and throughput of analysis pipelines. The steps in analyzing these images include registration, axon segmentation, soma detection, visualization and analysis of results. Several tools exist for these individual steps, but are rarely all part of an integrated pipeline and able to facilitate cloud-based collaboration [7, 9]. Further, many existing machine learning based tools are highly tuned to their training data and perform poorly when they encounter out-of-distribution artifacts or signal levels [6].

To address these challenges, we present BrainLine, an open-source, fully-integrated pipeline that performs registration, axon segmentation, soma detection, visualization, and analysis on whole-brain fluorescence volumes (Figure 1a). BrainLine combines state-of-the-art, already available open-source tools such as CloudReg [3] and ilastik [2] with brainlit, our Python package developed here. The BrainLine pipeline uses generalizable machine learning training schemes that adapt to out-of-distribution samples and facilitates cloud-based collaboration across institutions.

**Figure 1:**
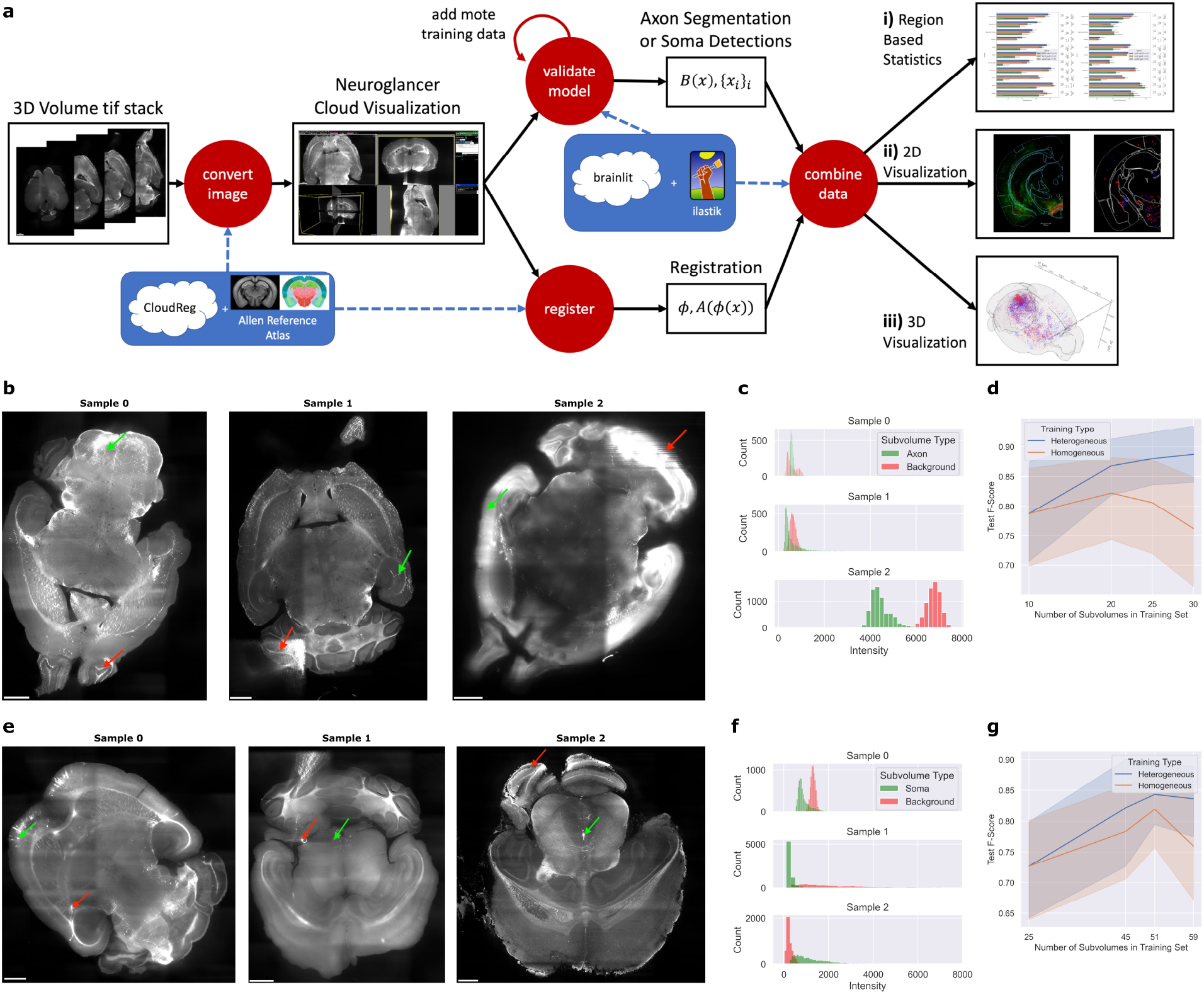
BrainLine allows for efficient processing of heterogeneous whole brain fluorescence volumes. **a** BrainLine combines CloudReg [3], ilastik [2] and our package, brainlit, to produce results in both quantitative (**a.i**) and visual (**a.ii-a.iii**) formats. **b** Example images with fluorescently labeled axon projections and arrows pointing to regions with (green) and without (red) labeled axons. **c** Intensity histograms of 20×20×20 voxel subvolumes located at the arrows in **b. d** Comparison between axon segmentation performance after training on subvolumes from different samples (heterogeneous) or the same sample (homogeneous). **e** Example images with fluorescently labeled cell bodies and arrows pointing to regions with (green) and without (red) labeled cell bodies. **f** Intensity histograms of 20×20×20 voxel subvolumes located at the arrows in **e. g** Comparison between soma detection performance after training on subvolumes from different brain samples (heterogeneous) or a single brain sample (homogeneous).

To share and interact with data across multiple institutions, BrainLine uses Amazon S3 to store data in precomputed format, so it can be viewed using Neuroglancer [1]. Specifically, we use CloudReg [3] for file conversion of the stitched image, and for image registration to the Allen atlas [10].

For axon segmentation and soma detection, we sought to leverage recent machine learning advances but experienced two major constraints. First, as generating ground truth image annotations is labor intensive, we wanted the approach to be effective on a small amount of training data. Second, images were provided to us in a sequential manner, and new samples would sometimes have unique artifacts or different levels of image quality (Figure 1b-c,e-f). We therefore sought a learning algorithm that could be quickly retrained on new data. Many learning algorithms assume that all training and testing data come from the same distribution and fail when this is not the case [8]. However, using our closed-loop training paradigm with ilastik [2], we were able to use a single ilastik project for all samples, only occasionally adding training data when difficult samples arose.

We used an ilastik pixel classification workflow for both axon segmentation and soma detection, but in the latter case we applied a size threshold to the connected components following segmentation. In both cases, the training approach was the same. For each new whole-brain volume, we identified a set of subvolumes (993 voxels for axons, 493 for somas) across a variety of brain regions, and annotated only a few slices (three for axons, five for somas) in each subvolume for our validation set. This strategy is similar to that employed in Friedmann et al. [5]. If our model could not achieve a satisfactory f-score on this validation dataset, we would annotate more subvolumes from the sample and add them to the training set until satisfactory performance was achieved.

We observed that this heterogeneous training procedure (i.e. training on multiple brain samples) often improved performance on other samples as well. In an experiment where we controlled the number of subvolumes used for training, this approach was at least as good as a homogeneous approach, where all training subvolumes came from a single brain sample (Figure 1d,g).

The pipeline can display the axon segmentation and soma detection results in a variety of ways, including brainregion-based bar charts accompanied by statistical tests (Fig. 1a.i), 2D plots with the atlas borders (Fig. 1a.ii), and 3D visualizations using brainrender (Fig. 1a.iii) [4]. Since every experimental design is unique, we designed our pipeline in a modular way, so investigators can pick and choose which components they want to incorporate in their own analyses. We also leverage existing software and file formats to facilitate interoperability [9].

BrainLine enables accelerated analysis of brain-wide connectivity through parallel programming, the use of cloud-compliant file formats, and a machine learning training scheme that generalizes across brain samples. As a result, BrainLine alleviates the need for investigators to build custom analysis pipelines from scratch, helping them characterize the morphology and connectivity profiles of neurons, and discover new neuronal subtypes. BrainLine is available as a set of thoroughly documented notebooks and scripts in our Python package brainlit: http://brainlit.neurodata.io/.

## Acknowledgements

This work is supported by NIH Grants RF1MH121539, U19AG033655, and RO1AG066184-01, NSF grants 2031985, 2014862 and the CAREER award. M.W. is supported by NIMH Grant K08MH113039. K.D. is supported by NIMH, NIDA, the NIH BRAIN Initiative, the Integrated Circuit Cracking NeuroNex Technology Hub funded by the National Science Foundation, the NOMIS Foundation, the Else Kröner Fresenius Foundation, the Gatsby Foundation and the AE Foundation.

